# Functionally dissecting claustrum along anterior-posterior axis in anxiety and higher sensitivity to cocaine

**DOI:** 10.1101/2022.12.30.522289

**Authors:** Ziheng Zhao, Zhaoyu Liu, Liying Chen, Wenwen Chen, Hao Guo, Jingjing Wang, Yuning Mai, Xiaoyan Wei, Jianhua Ding, Feifei Ge, Yu Fan, Xiaowei Guan

**Affiliations:** Department of Human Anatomy and Histoembryology, Nanjing University of Chinese Medicine, Nanjing, China; Department of Pharmacology, Nanjing Medical University, Nanjing, China

**Author notes:** These authors are equally contributed to this work (Ziheng Zhao, Zhaoyu Liu, Liying Chen). Correspondence to: Prof. Xiaowei Guan, Dr. Yu Fan.

## Abstract

Adolescent cocaine exposure (ACE) induce anxiety and higher sensitivity to drug addiction during adulthood. Here, we show that the claustrum is crucial for control of these two distinct psychiatric disorders in ACE mice. In the process of anxiety test, the CaMKII-positive neurons in median portion of claustrum (_M_Claustrum) were obviously triggered, and chemogenetic suppressing these neurons efficiently reduced ACE-induced anxiety in adulthood. While, the CaMKII-positive neurons in anterior portion of claustrum (_A_Claustrum) were obviously activated in response to cocaine-induced conditioned place preference (CPP), and chemogenetic suppressing these neurons efficiently blocked cocaine CPP in ACE mice during adulthood. Our findings dissociating specific sub-portions of claustrum for drug-related anxiety and susceptibility of addiction, extending our understanding to diverse functions of claustrum subregions.

## Introduction

Substance abuse in adolescents affects brain development and can influence health in adulthood, which is major health concern worldwide. Adolescent drug experience leads to the greater risk for developing addiction and/or anxiety, two prominent and distinct psychiatric disorders later in life (1–5), however, the price neuronal ensembles conferring vulnerability to these disorders remain unclear. The claustrum is a thin sheet of neuronal nucleus lying medially to insular cortex and laterally to putamen. Anatomically, the claustrum extensively connects with cerebral cortex in specific architecture of anterior-posterior and dorsal-ventral organization (6), implying different functions of sub-portions of claustrum along anterior-posterior axis. Recently, the claustrum shows greater activity in marijuana addicts (7, 8) and internet game addicts (9), implying a critical response of claustrum to addiction. One recent study shows that the claustrum mediates stress-induced anxiety in mice (10). The other recent study reported that claustrum, especially its dopamine receptor 1-containing (D1R^+^) neurons, is implicated in the acquisition and reinstatement of cocaine-preferred behaviors (11). These studies demonstrate that claustrum involves in both anxiety and addiction, however, little research literature is with respect to the specific sub-portion of claustrum for drug-related anxiety and addiction.

In the present study, basing on the fact that adolescent cocaine exposure (ACE) is critical risk factor for higher susceptibility of drug addiction and withdrawal anxiety later in life, we aims to dissociating specific sub-portions of claustrum for ACE-induced anxiety and higher sensitivity to drug addiction in mice during adulthood.

## Results and Discussion

In the present study, adolescent cocaine-exposed (ACE) male mice model was established by exposing male mice to cocaine from postnatal day 28 (P28) to P42, representing adolescent period in rodents (12). When growing up to adult age, mice were subjected to either elevated plus maze (EPM) or conditioned place preference (CPP) to evaluate withdrawal anxiety or addiction, respectively.

Although the claustrum has been investigated for hundreds of years, its precise function remains mysterious. Recent studies demonstrate that claustrum connects with the entire neocortical regions (6), and densely communicates with limbic structure and thalamus (13, 14), suggesting its potential role in emotion and cognition. The claustrum is a much narrow and long nucleus along anterior-posterior axis of brain, which could be divided into several subregions based on their different or specific projections with other brain area, suggesting diverse functions of claustrum subregions. Yet, few studies focus on the intrinsic fine-regional function of claustrum. Here, we separately observed the activating status of sub-portions of claustrum including anterior (_A_Claustrum), median (_M_Claustrum) and posterior (_P_Claustrum) in ACE mice during adulthood.

### The _M_Claustrum mediates ACE-induced anxiety in adulthood

To examine anxiety-like behaviors following ACE, mice were subjected to behavioral tests of EPM and open field test (OFT) in their adulthood (Figure 1A). When compared with adolescent saline exposure (ASE) mice, ACE mice exhibited less time spent in the open arms of EPM apparatus (Figure 1B) and in the central area of OFT box (Figure 1C) with similar total distances traveled in EPM apparatus (Figure 1B) and in the OFT box (Figure 1C). These results indicate that ACE induce anxiety without affecting their locomotor in mice during adulthood.

**Figure 1.**
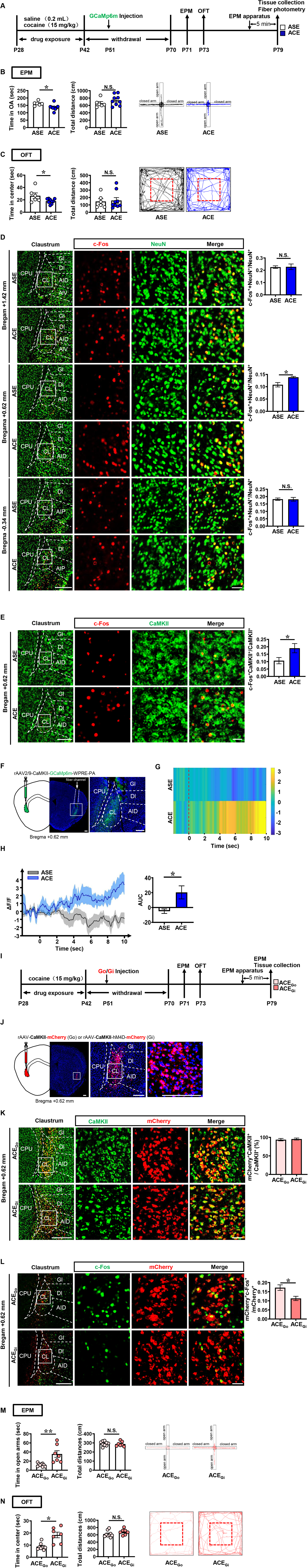
ACE enhances anxiety with evoked _M_Claustrum in adulthood. (**A**) Experimental timeline. (**B**) EPM test. Two-tailed unpaired t test. Left: t = 2.643, df = 13, *p* = 0.0203 versus ASE group; Right: t = 1.167, df = 13, *p* = 0.2643 versus ASE group. N.S., *p* > 0.05 versus ASE group. ASE group, n =6 animals; ACE group, n = 9 animals. (**C**) OFT. Two-tailed unpaired t test. Left: t = 2.181, df = 13, *p* = 0.0482 versus ASE group; Right: t = 0.2325, df = 13, *p* = 0.8197 versus ASE group. N.S., *p* > 0.05 versus ASE group. ASE group, n =6 animals; ACE group, n = 9 animals. (**D**) The percentage of c-Fos-positive neurons in claustrum. Two-tailed unpaired t test. Bregram +1.42 mm: t = 0.1102, df = 10, *p* = 0.9144 versus ASE group; Bregram +0.62 mm: t = 3.16, df = 10, *p* = 0.0102 versus ASE group; Bregram -0.34 mm: t = 0.07812, df = 10, *p* = 0.9393 versus ASE group. N.S., *p* > 0.05 versus ASE group. n =3 animals per group. Scale bar, 300 μm/50 μm. (**E**) The percentage of c-Fos-positive & CaMKII-positive neurons in _M_Claustrum. Two-tailed unpaired t test. Bregram +0.62 mm: t = 2.233, df = 10, *p* = 0.0496 versus ASE group. n =3 animals per group. Scale bar, 300 μm/50 μm. (**F**) Schematics and representative images of virus injection. Scale bar, 300 μm. (**G**) Heatmap of GCaMp6m fluorescence within the first 10 sec-duration in the EPM apparatus. (**H**) Quantification of ΔF/F and the AUC of ΔF/F. t = 2.718, df = 10, *p* = 0.0216 versus ASE group. n =6 animals per group. (**I**) Experimental timeline. (**J**) Schematics and representative images of virus injection. Scale bar, 300 μm. (**K**) Percentage of viral transfected neurons in CaMKII-positive neurons of _M_Claustrum. n = 3 animals per group. Scale bar, 300 μm/50 μm. (**L**) C-fos staining with mCherry fluorescence in _M_Claustrum. Two-tailed unpaired t test. t = 3.069, df = 10, *p* = 0.0119 versus ACE_Go_ group. n = 3 animals per group. Scale bar, 300 μm/50 μm. (**M**) EPM test. Two-tailed unpaired t test. Left: t = 3.678, df = 14, *p* = 0.0025 versus ACE_Go_ group; Right: t = 0.314, df = 14, *p* = 0.7582 versus ACE_Go_ group. N.S., *p* > 0.05 versus ACE_Go_ group. ACE_Go_ group, n =9 animals; ACE_Gi_ group, n = 7 animals. (**N**) OFT. Two-tailed unpaired t test. Left: t = 4.258, df = 14, *p* = 0.0008 versus ACE_Go_ group; Right: t = 1.414, df = 14, *p* = 0.1793 versus ACE_Go_ group. N.S., *p* > 0.05 versus ACE_Go_ group. ACE_Go_ group, n =9 animals; ACE_Gi_ group, n = 7 animals. ASE, adolescent saline exposure; ACE, adolescent cocaine exposure; ACE_Go_, control virus *rAAV-CaMKII-mCherry* injection to _M_Claustrum of adolescent cocaine-exposed mice; ACE_Gi_, *rAAV-CaMKII-hM4D (Gi)-mCherry* injection to _M_Claustrum of adolescent cocaine-exposed mice.

To map out the responsive portion of claustrum involved in anxiety, c-Fos, NeuN and CaMKII were used as markers to label neuronal activation, neuron and glutamatergic neuron, respectively. Compared with ASE mice, we found that _M_Claustrum neurons (Bregam: +0.62 mm, Figure 1D), but not _A_Claustrum (Bregam: +1.42 mm, Figure 1D) and _P_Claustrum (Bregam: -0.34 mm, Figure 1D), was activated following anxiety tests in ACE mice during adulthood, as indicated by the percentage of c-Fos-positive neurons in claustrum.

Since 90% neurons in claustrum belong to glutamatergic neurons (15, 16), we detect the activated CaMKII-positive neurons in _M_Claustrum of ACE mice. As shown in Figure 1E, we found that the percentage of c-Fos-positive neurons in _M_Claustrum CaMKII-positive neurons were higher following anxiety tests in ACE mice than those in ASE mice during adulthood.

To *in vivo* record real-time calcium signals of _M_Claustrum CaMKII-positive neurons during EPM test, *rAAV2/9-CaMKII-GCaMp6m-WPRE-PA* virus were injected into _M_Claustrum (Figure 1F). The “zero” point is set when mouse was put into EPM apparatus. Compared to ASE mice, the calcium signals in _M_Claustrum CaMKII-positive neurons were kept at higher levels within the first 10 sec-duration in the EPM apparatus (Figure 1G), and the area under curve (AUC) value of ΔF/F (Figure 1H) was higher in ACE mice than that in ASE mice during adulthood. These results indicate that the activation of _M_Claustrum, especially its CaMKII-positive neurons, be associated with the ACE-induced anxiety during adulthood.

To examine the role of ACE-activated _M_Claustrum CaMKII-positive neurons in anxiety, the designer receptors exclusively activated by designer drugs (DREADDs)-inhibiting CaMKII-positive neurons were performed by bilaterally infusing *rAAV-CaMKII-hM4D-mCherry* (ACE_Gi_ group) into _M_Claustrum. The *rAAV-CaMKII-mCherry* (ACE_Go_ group) was used as controls (Figure 1I, 1J). Above 90% CaMKII-positive neurons within _M_Claustrum were transfected with virus (Figure 1K). Compared with ACE_Go_ mice, clozapine N-oxide (CNO) treatment attenuated activation of CaMKII-positive neurons in _M_Claustrum, as indicated by lower percentage of c-Fos-positive neurons in transfected CaMKII-positive neurons in _M_Claustrum of ACE_Gi_ mice (Figure 1L). Following CNO treatment, ACE_Gi_ mice spent more time in the open arms of EPM apparatus (Figure 1M) and in the central area of the OFT box (Figure 1N) with similar distance traveled in the EPM apparatus (Figure 1M) and in OFT box (Figure 1N), when compared to ACE_Go_ controls.

Recently, Niu et al. (10) report that activating claustrum elicit stress-related anxiety, indicating a potential role of claustrum in anxiety. Here, using ACE-induced anxiety models, we found that adolescent cocaine exposure specifically triggers _M_Claustrum CaMKII-positive neurons in adulthood, and chemogenetic suppressing these neurons efficiently reduced anxiety in ACE mice during adulthood. These results demonstrate that _M_Claustrum subregion involves in coding drug-related anxiety.

### The _A_Claustrum mediates ACE-induced higher sensitivity to cocaine addiction in adulthood

Mice were subjected to CPP to evaluate the sensitivity to cocaine in adulthood. Here, 1 mg/kg of cocaine (insufficient to produce CPP normally) was used to induce CPP in adult mice (Figure 2A). As shown in Figure 2B-D, ASE mice did not acquire cocaine CPP during adulthood (ASE-C) following 4-day cocaine CPP training, while ACE mice acquired cocaine CPP during adulthood (ACE-C). The ΔCPP score was higher in ACE-C mice than that in ASE-C mice. These results indicate that adolescent cocaine exposure induce higher sensitivity to cocaine during adulthood, which present more vulnerability to drug addiction.

**Figure 2.**
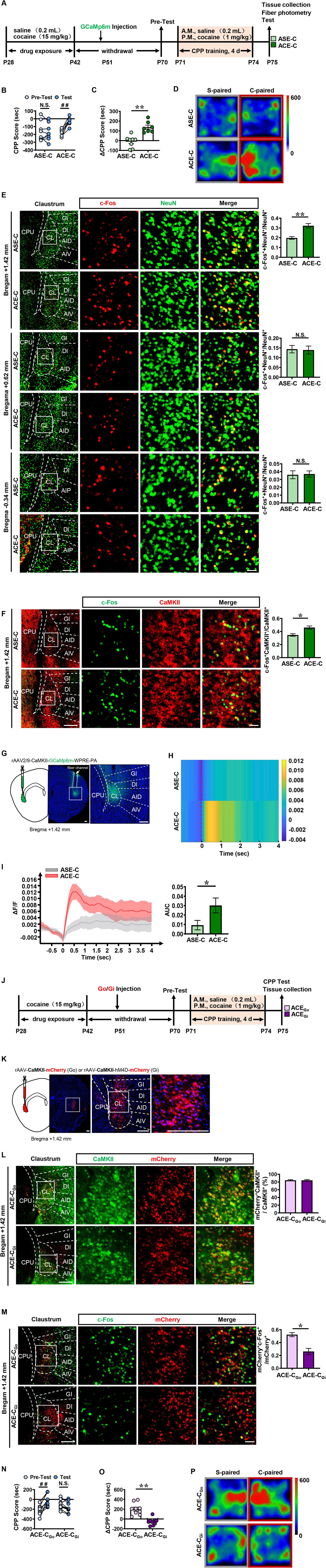
ACE increases higher sensitivity to cocaine addiction with evoked _A_Claustrum in adulthood. (**A**) Experimental timeline. (**B**) CPP score during the pre-test and CPP test. Two-way ANOVA with Sidak’s multiple comparisons test. ASE-C: t = 0.5009, df = 13, *p* = 0.8593 versus Pretest of ASE-C group; ACE-C: t = 6.220 df = 13, ##, *p* < 0.0001 versus Pre-test of ACE-C group. N.S., *p* > 0.05 versus Pre-test of ASE-C group. ASE-C group, n = 8 animals; ACE-C group, n = 7 animals. (**C**) ΔCPP score (CPP test minus pre-test) of the two group. Two-tailed unpaired t test. t = 4.884, df = 13, *p* = 0.0003 versus ASE-C group. ASE-C group, n =8 animals; ACE-C group, n = 7 animals. (**D**) Heatmap of spent duration by mice in CPP apparatus. (**E**) The percentage of c-Fos-positive neurons in claustrum. Two-tailed unpaired t test. Bregram +1.42 mm: t = 4.628, df = 10, *p* = 0.0009 versus ASE-C group; Bregram +0.62 mm: t = 0.1565, df = 10, *p* = 0.8788 versus ASE-C group; Bregram -0.34 mm: t = 0.1469, df = 10, *p* = 0.8862 versus ASE-C group. N.S., *p* > 0.05 versus ASE-C group. n =3 animals per group. Scale bar, 300 μm/50 μm. (**F**) The percentage of c-Fos-positive & CaMKII-positive neurons in _A_Claustrum. Two-tailed unpaired t test. Bregram +1.42 mm: t = 3.803, df = 4, *p* = 0.0191 versus ASE-C group. n =3 animals per group. Scale bar, 300 μm/50 μm. (**G**) Schematics and representative images of virus injection. Scale bar, 300 μm. (**H**) Heatmap of GCaMp6m fluorescence at first 4 sec when mice entering cocaine-paired chamber. (**I**) Quantification of ΔF/F and the AUC of ΔF/F. t = 2.523, df = 10, *p* = 0.0499 versus ASE-C group. n =6 animals per group. (**J**) Experimental timeline. (**K**) Schematics and representative images of virus injection. Scale bar, 300 μm. (**L**) Percentage of viral transfected neurons in CaMKII-positive neurons of _A_Claustrum. n =3 animals per group. Scale bar, 300 μm/50 μm. (**M**) C-fos staining with mCherry fluorescence in _A_Claustrum. Two-tailed unpaired t test. t = 4.528, df = 4, *p* = 0.0106 versus ACE-C_Go_ group. n =3 animals per group. Scale bar, 300 μm/50 μm. (**N**) CPP score during the pre-test and CPP test. Two-way ANOVA with Sidak’s multiple comparisons test. ACE-C_Go_: t = 6.53, df = 17, ##,*p* < 0.0001 versus Pretest of ACE-C_Go_ group; ACE-C_Gi_: t = 1.937 df = 17, *p* = 0.1343 versus Pre-test of ACE-C_Gi_ group. N.S., *p* > 0.05 versus Pre-test of ACE-C_Gi_ group. ACE-C_Go_ group, n =9 animals; ACE-C_Gi_ group, n = 10 animals. (**O**) ΔCPP score (CPP test minus pre-test) of the two group. Two-tailed unpaired t test. t = 5.836, df = 17, *p* < 0.0001 versus ACE-C_Go_. ACE-C_Go_ group, n =9 animals; ACE-CGi group, n = 10 animals. (**P**) Heatmap of spent duration by mice in CPP apparatus. ASE-C, subthreshold-dosed administration of cocaine in adolescent saline-exposed mice; ACE-C, subthreshold-dosed administration of cocaine in adolescent cocaine-exposed mice; ACE-C_Go_, control virus *rAAV-CaMKII-mCherry* injection to _A_Claustrum of ACE-C mice; ACE-C_Gi_, *rAAV-CaMKII-hM4D (Gi)-mCherry* injection to _A_Claustrum of ACE-C mice.

Compared to ASE-C mice, _A_Claustrum neurons (Bregam: +1.42 mm, Figure 2E), but not _M_Claustrum (Bregam: +0.62 mm, Figure 2E) and _P_Claustrum (Bregam: -0.34 mm, Figure 2E) neurons of ACE-C mice were more activated in response to cocaine CPP during adulthood, as indicated by the percentage of c-Fos-positive neurons in claustrum (Figure 2E). As shown in Figure 2F, the percentage of c-Fos-positive neurons in _A_Claustrum CaMKII-positive neurons were higher following cocaine-induced CPP in ACE-C mice than those in ASE-C mice during adulthood. The “zero” point for *in vivo* recording *GCaMp6m* is identified as the beginning of mouse entering into cocaine-paired chamber during CPP test. Compared to ASE-C mice, the calcium signals in _A_Claustrum CaMKII-positive neurons were kept at higher levels in first 4-sec recording phase (Figure 2G, 2H), and the AUC value of ΔF/F was higher in ACE-C mice than that in ASE-C mice during adulthood (Figure 2I). These results indicate that the activation of _A_Claustrum, especially its CaMKII-positive neurons, be associated with the ACE-induced higher sensitivity to cocaine during adulthood.

The *rAAV-CaMKII-hM4D-mCherry* (ACE-C_Gi_ group) were bilaterally injected into _A_Claustrum, and *rAAV-CaMKII-mCherry* (ACE-C_Go_ group) was used as controls (Figure 2J, 2K). Above 80% CaMKII-positive neurons within _A_Claustrum were transfected with virus (Figure 2L). Compared with ACE-C_Go_ mice, CNO treatment suppressed CaMKII-positive neurons in _A_Claustrum, as indicated by lower percentage of c-Fos-positive neurons in transfected CaMKII-positive neurons in _A_Claustrum of ACE-C_Gi_ mice (Figure 2M). Following CNO treatment, ACE-C_Go_ mice still exhibited cocaine-induced CPP during adulthood (Figure 2N). While, CNO treatment efficiently blocked cocaine-induced CPP in ACE-C_Gi_ mice during adulthood (Figure 2N). The ΔCPP score in ACE-C_Gi_ mice was much lower than that in ACE-C_Go_ mice (Figure 2O, 2P).

Recently, Terem et al. (11) found that D1R^+^ neurons in claustrum, especially those connected with prefrontal cortex, mediate the acquisition and reinstatement of cocaine-related CPP. Here, using models of ACE-induced higher sensitivity to cocaine in adulthood, we found that _A_Claustrum, especially its CaMKII-positive neurons, were obviously more activated in response to cocaine-induced CPP, and chemogenetic suppressing these neurons efficiently blocked cocaine-preferred behavior in ACE mice during adulthood. These results demonstrate that _A_Claustrum subregion involves in coding susceptibility of substance abuse.

In conclusion, our findings reveal that adolescent cocaine exposure-induced anxiety and higher sensitivity to cocaine recruit different portions of claustrum (median and anterior, respectively), extending our understanding to diverse functions of claustrum subregions (Figure 3).

**Figure 3.**
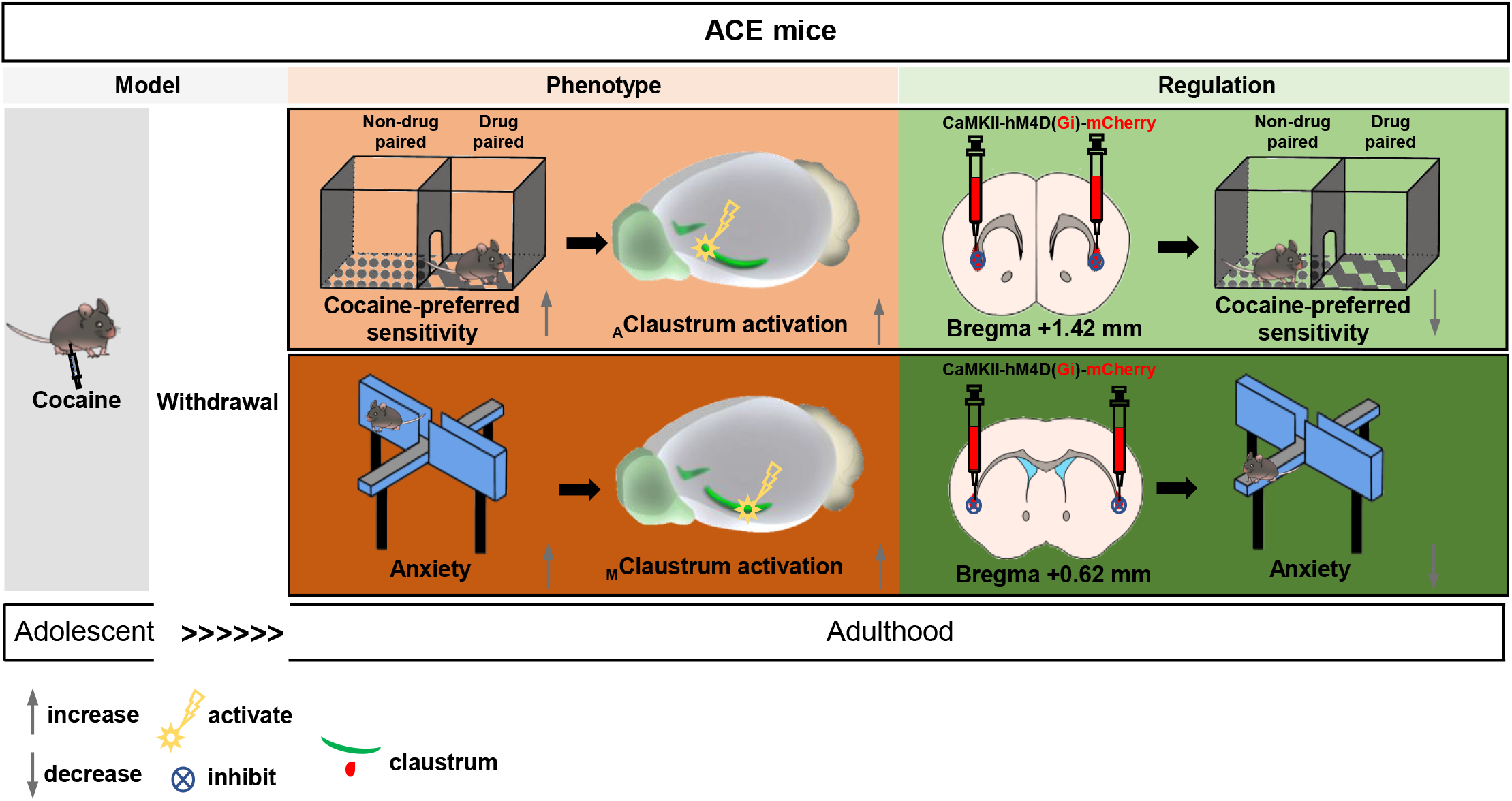
Schematic summary of the present study. Adolescent cocaine-exposed male mice (ACE) models were established by administrating cocaine of 15 mg/kg once daily during adolescent period. When growing to adult age, mice were subjected to either elevated plus maze (EPM) or conditioned place preference (CPP) to evaluate anxiety and addiction, respectively. The CaMKII-positive neurons in median portion of claustrum (_M_Claustrum) were obviously triggered in the process of anxiety test, and chemogenetic suppressing these neurons efficiently reduced ACE-induced anxiety in adulthood. While, the CaMKII-positive neurons in anterior portion of claustrum (_A_Claustrum) were obviously activated in response to subthreshold dose of cocaine-induced conditioned place preference (CPP), and chemogenetic suppressing these neurons efficiently blocked cocaine CPP in ACE mice during adulthood.

## Materials and methods

### Animals

Male C57BL/6 wild type (WT) mice weighing 12-16 g were used. All animals were housed at constant humidity (40~60%) and temperature (24 ± 2°C) with a 12-hour light/dark cycle (lights on at 8 a.m.) and allowed free access to food and water. Handling of the animals was carried out for three days before onset of experiments. All procedures were carried out in accordance with the National Institutes of Health Guide for the Care and Use of Laboratory Animals and approved by the Institutional Animal Care and Use Committee (IACUC) at Nanjing University of Chinese Medicine.

### Drug treatment

From P28 to P42, male C57BL/6 WT mice were assigned to receive a single daily injection of cocaine hydrochloride (15 mg/kg, dissolved in saline, i.p., Qinghai Pharmaceutical, China) or saline (0.2 mL, i.p.) once daily at 10 a.m. for 15 consecutive days. From P42-P70, these mice were kept at their home cage (four mice per cage). From P71 to P73, the cohort 1 and 2 of mice were subjected to the related anxiety-like behavioral tests, the mice were sacrificed after stimulus of EPM procedure on P79. The cohort 3 and 4 of mice were subjected to CPP training by subthreshold dose of cocaine hydrochloride (1 mg/kg) and test on P75. The brain tissue was collected for immunofluorescence.

### Behavioral tests

All behavioral tests were conducted during light cycle. In the current study, two cohorts of the adolescent cocaine exposure mice were used to assess anxiety-like behaviors, two cohorts were subjected to CPP. All data were recorded and analyzed by TopScan H system (TopScan, USA). The mice were kept at their home cage until experiments.

EPM and OFT were performed to assess anxiety-like behaviors in cohort 1 and 2 of the ACE mice on P71 and P73, respectively. CPP training was performed in cohort 3 and 4 of the ACE mice from P71 to P74. The EPM apparatus (TopScan, Kansas City, MO, USA) consists of 4 elevated arms (52 cm above the floor) arranged in a cross pattern with two 1-cm walled arms (open arms) and two 40-cm walled arms (closed arms). Each mouse was placed in the center portion of the EPM with its head facing an open arm and allowed freely to explore the maze for 5 min. The time spent in and the entries into the open arms and the total distance traveled in the apparatus were recorded. In OFT, mice were placed gently in the center of the open field apparatus (50 × 50 × 50 cm) individually, and the time spent in and the entries into the central area (16× 16 × 16 cm) of the box and the total distance traveled in the box were recorded for 5 min with a camera positioned overhead.

CPP were performed to evaluate the sensitivity to cocaine-induced preference in mice. The CPP was performed in the TopScan3D CPP apparatus (CleverSys, VA, USA), which is constructed of two distinct chambers (15 × 15 × 23 cm each) separated by a removable guillotine door. The CPP procedure consisted of three phases: the pre-conditioning test (Pre-test, P70), conditioning (CPP training, P71-74) and post-conditioning test (Test, P75). Mice were allowed a 15-min free exploration to the two chambers as the baseline preference, then equally divided into two sub-groups. Here, subthreshold doses of cocaine (1 mg/kg), which is believed not enough to induce CPP in naïve mouse, were used to induce CPP. During CPP training, mice were injected with saline (0.2 mL, i.p.) in the morning and cocaine (1 mg/kg, i.p.) in the afternoon once daily. After each injection, the mice were confined to one chamber (non-drug-paired chamber or drug-paired chamber) for 45 min. During the Pre-test and Test, mice freely access two chambers for 15 min. The CPP score is the time spent in drug-paired chamber minus that in non-drug-paired chamber, and the ΔCPP score is the test CPP score minus the pre-test CPP score.

### Immunofluorescence

Mice brains were collected within 90 min after EPM test or CPP test and perfused with 4% paraformaldehyde (PFA). The coronal brain sections (30 μm) were cut on a cryostat (Leica, Germany). The sections were incubated in 0.3% (v/v) Triton X-100 for 0.5 h, blocked with 5% donkey serum for 1.5 h at room temperature, and incubated overnight at 4°C with the following primary antibodies: rabbit anti-c-Fos (1:2000, RRID: AB_2247211, Cell Signalling Technology, USA), mouse anti-NeuN (1:800, RRID: AB_2298772, Millipore, USA), and mouse anti-CaMKIIα (1: 100, RRID: AB_626789, Santa Cruz, USA), followed by the corresponding fluorophore-conjugated secondary antibodies for 1 h at room temperature. The following secondary antibodies were used here: Alexa Fluor 555-labeled donkey anti-rabbit secondary antibody (1:500, RRID: AB_162543, Invitrogen, USA), Alexa Fluor 488-labeled donkey anti-mouse secondary antibody (1:500, RRID: AB_141607, Invitrogen, USA). Fluorescence signals were visualized using a Leica DM6B upright digital research microscope (Leica, Germany) or Leica Dmi8 (Leica, Germany).

### Fiber photometry

On P51, mice were anesthetized with 2% isoflurane in oxygen, and were fixed in a stereotactic frame (RWD, China). A heating pad was used to maintain the core body temperature of the animals at 36°C. The coordinates were defined as dorso-ventral (DV) from the brain surface, anterior-posterior (AP) from bregma and medio-lateral (ML) from the midline (in mm). A volume of 70 nL of the *rAAV2/9-CaMKII-GCaMp6m* (PT-0111, 3.24E+12 vg/ml, Brain VTA, China) virus was bilaterally injected into _M_Claustrum (AP + 0.62 mm, ML ± 2.80 mm, DV – 3.50 mm) or _A_Claustrum (AP + 1.42 mm, ML ± 3.05 mm, DV – 3.50 mm) at a rate of 10 nL/min. In Experiment 1, an optical fiber (200 μm outer diameter, 0.37 numerical aperture (NA), AOGUAN Biotech, China) was placed 100 μm above the viral injection site. The calcium-dependent fluorescence signals were obtained by stimulating cells that transfected GCaMp6m virus with a 470 nm LED (35-40 μW at fiber tip), while calcium-independent signals were obtained by stimulating these cells with a 405 nm LED (15-20 μW at fiber tip). The LED lights of 470 nm and 405 nm were alternated at 40 fps and light emission was recorded using sCMOS camera containing the entire fiber bundle (2 m in length, NA = 0.37, 200 μm core, Thinkerbiotech, China). The analog voltage signals fluorescent was filtered at 30 Hz and digitalized at 100 Hz. The GCaMp6m signals were recorded and analyzed by ThinkerTech TrippleColor MultiChannel fiber photometry Acquisition Software and ThinkerTech TrippleColor MultiChannel fiber photometry Analysis Package (Thinkerbiotech, China, China), respectively. F0 is the baseline fluorescence signal which was recorded for 2 sec before mice entering into the EPM. F is the real-time fluorescence signal which was recorded for 10 sec after mice entering into the EPM. Each recording trail was with 12 sec recording duration. In Experiment 2, an optical fiber (200 μm outer diameter, 0.37 NA, Inper Ltd., China) was placed 100 μm above the viral injection site. The calcium-dependent fluorescence signals were obtained by stimulating cells that transfected GCaMp6m virus with a 470 nm LED (35-40 μW at fiber tip), while calcium-independent signals were obtained by stimulating these cells with a 405 nm LED (15-20 μW at fiber tip). The LED lights of 470 nm and 405 nm were alternated at 66 fps and light emission was recorded using sCMOS camera containing the entire fiber bundle (2 m in length, NA = 0.37, 200 μm core, Inper Ltd.). The analog voltage signals fluorescent was filtered at 30 Hz and digitalized at 100 Hz. The GCaMp6m signals were recorded and analyzed by Inper Studio Multi Color EVAL15 software (Inper Ltd., China) and Inper Data Process V0.5.9 (Inper Ltd., China), respectively. F0 is the baseline fluorescence signal which was recorded for 1 sec before mice entering into the cocaine-paired chamber. F is the real-time fluorescence signal which was recorded for 4 sec after mice entering into the cocaine-paired chamber during CPP test. Each recording trail was with 5 sec recording duration. The values of ΔF/F are calculated by (F-F0)/F0. The area under curve (AUC) is the integral under recording duration related to corresponding baseline at every trial.

### Designer receptors exclusively activated by designer drugs

On P51, mice were anesthetized with 2% isoflurane in oxygen, and were fixed in a stereotactic frame (RWD, China). A heating pad was used to maintain the core body temperature of the animals at 36°C. A volume of 70 nL of *rAAV-CaMKII-hM4D(Gi)-mCherry* or *rAAV-CaMKII-mCherry(Go)* bilaterally into the _M_Claustrum (AP + 0.62 mm, ML ± 2.80 mm, DV – 3.50 mm) or _A_Claustrum (AP + 1.42 mm, ML ± 3.05 mm, DV – 3.50 mm) at a rate of 10 nL/min. After surgery, mice were maintained at home cage about 3 weeks. Mice were injected with CNO (1 mg/kg, i.p., MCE Co, USA) 30 min before all behavioral tests.

### Statistical analysis

Statistical analysis was carried out using GraphPad Prism 8.0.2 software. All data are presented as the Mean ± SEM. The data of CPP score are analyzed by two-way analysis of variance (ANOVA), while CPP score between Pre-test and Test in each mouse were analyzed by paired t-tests. Other data were analyzed by unpaired t-tests. All statistical significance was set as *p* < 0.05.

## Author contributions

Zhao Z, Liu Z and Chen L have equal contribution to the manuscript. Zhao Z, Liu Z and Chen L performed behavioral tests, morphological tests and virus-related experiment. Chen W, Guo H, Wang J, Mai Y, Wei X and Ding J assist the experiment and data analysis. Fan Y and Ge F performed data analysis. Guan X and Fan Y wrote the manuscript. Guan X developed the overall concept.

## Acknowledgments

This work is supported by National Natural Science Foundation of China (82271531 and 82071495) and Natural Science Foundation of the Higher Education Institutions of Jiangsu Province, China (21KJB360007).

## Conflict of interests

The authors have no conflict of interests to declare.

## Data availability statement

The data that support the findings of this study are available from the corresponding author upon reasonable request.

## Notes

### Competing Interest Statement

The authors have declared no competing interest.

### Summary of Updates

The manuscript has been revised.

